# Applying Spatial Statistics to Spatial Transcriptomics Reveals Local Association Between M2-like Macrophages and Fibrosis in Diabetic Kidney Disease

**DOI:** 10.64898/2026.05.31.729068

**Authors:** Kanako Terakawa, Hideaki Kawaguchi, Masaomi Nangaku, Imari Mimura

## Abstract

Renal fibrosis is the common final pathway of chronic kidney disease (CKD), driven in part by myofibroblast-mediated extracellular matrix deposition. M2 macrophages, hereafter referred to as MAC-M2, have been implicated in renal fibrosis, yet whether M2 macrophages are pro- or anti-fibrotic remains controversial, and the spatial context in which MAC-M2-fibrosis coupling occurs is unknown. Here, we applied geographically weighted regression (GWR), a spatial statistical method, to Visium spatial transcriptomics data from diabetic kidney disease (DKD) to characterize spatially resolved high-coupling spots where MAC-M2-fibrosis coupling is significantly positive. In a small DKD cohort (n=6), GWR identified high-coupling spots enriched for B cell and tertiary lymphoid structure (TLS)-like immune signatures, suggesting that the GWR-defined regions captured biologically meaningful immune microenvironments. To gain statistical power for differential gene expression (DEG) analysis, we then applied the same pipeline to the larger Kidney Precision Medicine Project (KPMP) DKD cohort (n=30), in which high-coupling spots showed upregulation of IgE-related immune genes *(IGHE*, *FCER1A*) together with the mast cell tryptase *TPSB2*. These findings suggest that IgE-related immune responses may be present within DKD fibrotic microenvironments characterized by local MAC-M2-fibrosis coupling. As a disease comparison, we further applied the pipeline to a KPMP hypertensive kidney disease (HKD) cohort (n = 27), where high-coupling spot signatures were distinct from DKD and did not show enrichment of IgE-related genes. Together, this study provides the first application of GWR to kidney spatial transcriptomics and suggests that IgE-related immune responses may be a feature of DKD fibrotic microenvironments in which M2 macrophages are locally associated with fibrosis.

**Highlights:**

- Geographically weighted regression (GWR) maps spatially variable M2 macrophage-fibrosis coupling in diabetic kidney disease (DKD).
- GWR-defined high-coupling spots show immune activation and loss of kidney-specific programs.
- The GWR-based analysis was replicated across two independent DKD cohorts.
- *IGHE*, *FCER1A*, and the mast cell marker *TPSB2* are enriched in high-coupling spots in the KPMP DKD cohort.

## Introduction

Diabetic kidney disease (DKD) is the leading cause of kidney failure with replacement therapy worldwide, affecting approximately 40% of patients with diabetes and accounting for over half of all cases of end-stage kidney disease globally^1^. DKD is characterized by spatially heterogeneous tissue injury: within a single kidney biopsy, regions of varying degrees of fibrosis, inflammation, tubular atrophy, and glomerulosclerosis coexist in a complex mosaic, reflecting the interplay between adaptive and maladaptive repair mechanisms across different tissue compartments^2^. This spatial heterogeneity has important implications for understanding disease pathogenesis, as global or bulk-tissue analyses inevitably average across microenvironments with distinct biological states, potentially obscuring the cellular interactions that drive progressive injury.

The kidney is among the most cellularly complex organs, containing highly specialized epithelial, endothelial, interstitial, and immune cell populations distributed across distinct nephron segments and tissue compartments^3^. The kidney also exhibits stereotyped tissue-level architecture, with strong polarity along the cortical-medullary axis and along individual nephron segments, such that each cell type occupies a defined spatial niche. This combination of extensive cellular heterogeneity and strong spatial polarity makes the kidney a natural target for spatial statistical analysis.

Renal fibrosis is the common final pathway of CKD progression, driven by myofibroblast-mediated extracellular matrix (ECM) deposition. Fibrotic remodeling involves activation of stromal populations, including resident fibroblasts and perivascular cells, which contribute to matrix-producing myofibroblasts^2,4^. Macrophages, in turn, can modulate fibrogenesis through cytokine- and growth factor-mediated interactions with stromal, epithelial, and immune compartments^5^. However, the net role of M2 macrophages in kidney fibrosis remains controversial. On one hand, M2-polarized macrophages express anti-inflammatory cytokines (IL-10, IL-4) and growth factors (TGF-β, PDGF) that can promote both tissue repair and ECM deposition. On the other hand, adoptive transfer and depletion studies in mice have produced conflicting results regarding whether M2 macrophages accelerate or attenuate renal fibrosis, likely reflecting context-dependence of M2 macrophage function^6^. A key unanswered question is whether the association between M2 macrophages and fibrosis is spatially uniform across the kidney or concentrated within specific pathological microenvironments. Addressing this question requires an analytical framework that can quantify local associations at the level of individual tissue spots.

Spatial transcriptomics enables gene expression profiling while preserving tissue architecture, offering a key advantage over bulk RNA-seq and single-cell RNA-seq for the study of spatially heterogeneous disease. In the kidney, recent studies have identified distinct fibrotic, immune, tubular, and glomerular microenvironments using spatial transcriptomic approaches^7,8^. However, current analytical frameworks are still largely descriptive. Co-localization analyses can identify which cell types or signals appear in proximity, and microenvironment classification approaches can group spots into discrete categories, but these methods do not directly quantify where the local relationship between two continuous biological variables becomes stronger or weaker across tissue space. As a result, spatial transcriptomics has not yet fully realized its potential for identifying spatially restricted disease-associated relationships.

Addressing this gap requires integration of spatial transcriptomics with quantitative methods developed outside the biomedical field. Zormpas et al.^9^ proposed that spatial statistical methods originally developed in geography, including geographically weighted regression (GWR), can be applied to spatial transcriptomics data. In conventional regression, sometimes called global regression, the relationship between two variables is summarized by a single coefficient assumed to hold uniformly across the entire tissue. By contrast, GWR is a local regression method: it estimates a separate relationship at each spatial location by weighting nearby spots more strongly than distant ones, yielding a location-specific coefficient for every spot. This distinction between global and local modeling is central to the present approach. This framework is therefore well suited to detect spatial non-stationarity and to identify regions where two biological variables are more tightly coupled than in surrounding tissue. In the context of DKD, such an approach provides a way to move beyond descriptive spatial mapping and instead quantify where M2 macrophage abundance is locally associated with fibrosis. Despite this conceptual promise, GWR has not yet been applied to kidney spatial transcriptomics.

Here, we present the first application of GWR to kidney spatial transcriptomics, with a focus on DKD. We first applied this framework to a small previously published DKD cohort (n=6; Abedini et al.^7^) as a proof-of-concept. To increase statistical power for downstream differential expression analysis, we then applied the same pipeline to the larger KPMP DKD cohort (n=30) as our primary analysis. Finally, to assess whether the observed transcriptional features were specific to DKD rather than a general property of chronic fibrotic kidney disease, we applied the identical pipeline to the KPMP hypertensive kidney disease (HKD) cohort (n=27). Using this interdisciplinary approach integrating nephrology, spatial transcriptomics, and spatial statistical modeling, we sought to identify spatially restricted regions of MAC-M2-fibrosis coupling and to define the molecular features associated with these regions in DKD.

## Results

### Overview

To identify spatially restricted regions of MAC-M2-fibrosis coupling within kidney tissue, we developed a GWR-based analysis pipeline for Visium spatial transcriptomics data (Figure 1A, B). Per-spot M2 macrophage abundance was estimated by label transfer from a kidney single-cell reference atlas^8^, and a fibrosis score was derived from myofibroblast marker genes. GWR was fitted independently to each sample to obtain spot-level MAC-M2-fibrosis regression coefficients, and spots with a significantly positive coefficient (Benjamini-Hochberg (BH)-adjusted p < 0.05) were defined as high-coupling spots; among spots with evaluable GWR estimates, all remaining spots were used as the reference group. We applied this pipeline sequentially to three cohorts (Figure 1C): a small DKD dataset (n=6; Abedini et al.^7^) as initial proof-of-concept, the larger KPMP DKD cohort (n=30) as our higher-powered primary analysis, and the KPMP HKD cohort (n=27) for testing disease specificity. All six samples in the proof-of-concept cohort passed sample-level quality control, while one KPMP DKD sample and four KPMP HKD samples were excluded by quality control (QC), leaving 6, 29, and 23 samples respectively for GWR analysis. Of these, 3, 14, and 9 samples reached the threshold for downstream pseudobulk DEG analysis (at least 20 high-coupling spots and at least 20 reference spots; Figure 1C). Clinical characteristics of the KPMP DKD and HKD cohorts are summarized in Table 1. As expected, the two cohorts differed in several baseline clinical variables, including age, baseline eGFR, glycemic status, and diabetes-related history. Detailed methodology is provided in STAR Methods.

**Figure 1.**
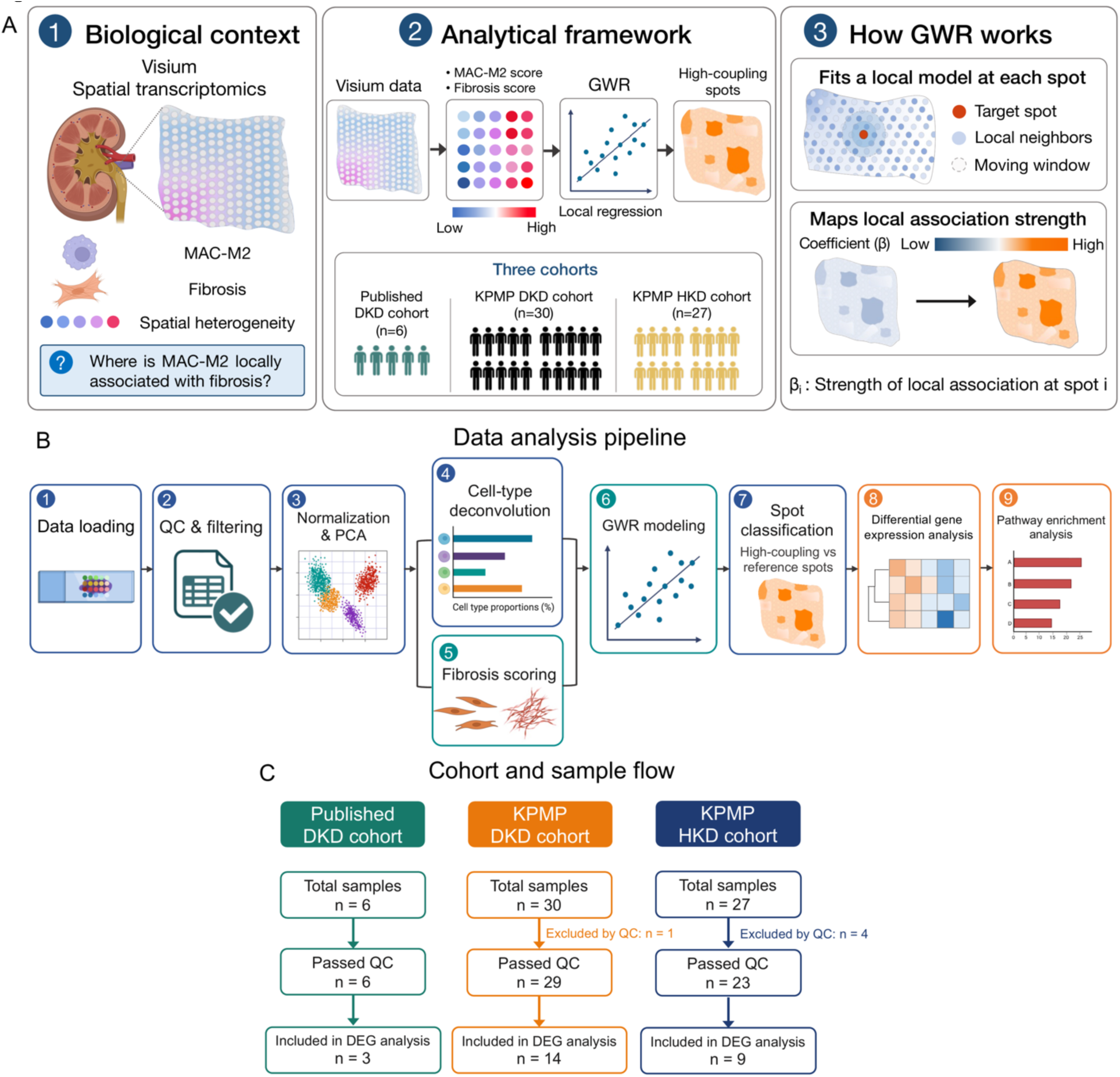
Study concept, analysis pipeline, and cohort overview. (A) Conceptual framework. Left, the biological motivation : DKD tissue exhibits spatial heterogeneity in both M2 macrophage abundance and fibrosis, and the question addressed is where the local association between M2 macrophages and fibrosis is strongest. Middle, analytical strategy : Visium spatial transcriptomics is processed to derive a per-spot M2 macrophage score and fibrosis score, GWR is applied per sample to estimate a local MAC-M2-fibrosis regression coefficient at each spot, and spots with significantly positive coefficients are classified as high-coupling spots. Right, GWR operation : at each target spot, a moving-window local regression is fitted using neighboring spots within an adaptive bandwidth, yielding a spot-level coefficient β that maps the strength of the local MAC-M2-fibrosis association across the tissue. (B) Step-by-step analytical pipeline: (1) data loading, (2) sample- and spot-level quality control (median nFeature ≥ 500, median mitochondrial percentage ≤ 25%, minimum 100 spots per sample), (3) normalization and PCA, (4) spot-level cell-type deconvolution by label transfer from the KPMP single-cell RNA-seq reference atlas^8^, (5) per-spot fibrosis scoring using AddModuleScore with myofibroblast marker genes, (6) GWR modeling (Fibrosis Score ∼ M2 fraction + log10 UMI, adaptive bandwidth, bisquare kernel, AIC-selected), (7) spot classification into high-coupling and reference spots (BH-adjusted p < 0.05 on positive M2 coefficient), (8) pseudobulk differential gene expression analysis (edgeR, paired design ∼SampleID + Condition) on samples with ≥ 20 high-coupling and ≥ 20 reference spots, and (9) pathway enrichment analysis (GO biological process, GSEA Hallmark). (C) Cohort flow diagram for the three cohorts analyzed: a previously published DKD dataset (n=6, proof-of-concept^7^), the KPMP DKD cohort (n=30, primary analysis), and the KPMP HKD cohort (n=27, disease-specificity test). The diagram shows total samples, samples passing QC (DKD n=6 → 6; KPMP DKD n=30 → 29; KPMP HKD n=27 → 23), and samples included in DEG analysis after meeting the ≥ 20-spots threshold for both classes (DKD n=3; KPMP DKD n=14; KPMP HKD n=9). Created in BioRender. Terakawa, K. (2026) https://BioRender.com/cjnt04w

**Table 1.**
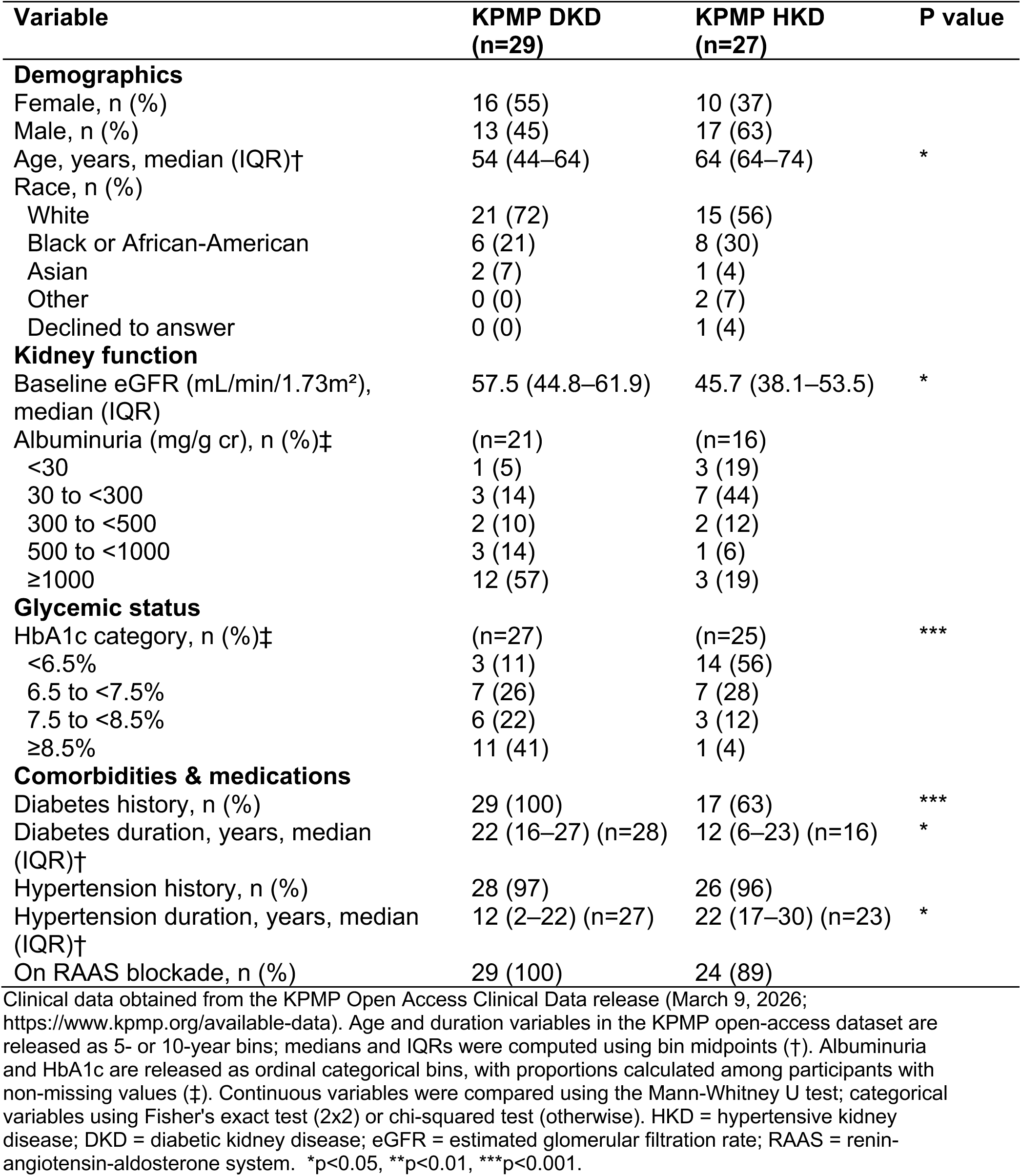
Clinical characteristics of KPMP DKD and HKD cohorts.

| Variable | KPMP DKD<br>(n=29) | KPMP HKD<br>(n=27) | P value |
| --- | --- | --- | --- |
| **Demographics** |  |  |  |
| Female, n (%) | 16 (55) | 10 (37) |  |
| Male, n (%) | 13 (45) | 17 (63) |  |
| Age, years, median (IQR)† | 54 (44–64) | 64 (64–74) | * |
| Race, n (%) |  |  |  |
| White | 21 (72) | 15 (56) |  |
| Black or African-American | 6 (21) | 8 (30) |  |
| Asian | 2 (7) | 1 (4) |  |
| Other | 0 (0) | 2 (7) |  |
| Declined to answer | 0 (0) | 1 (4) |  |
| **Kidney function** |  |  |  |
| Baseline eGFR (mL/min/1.73m<sup>2</sup>), median (IQR) | 57.5 (44.8–61.9) | 45.7 (38.1–53.5) | * |
| Albuminuria (mg/g cr), n (%)‡ | (n=21) | (n=16) |  |
| <30 | 1 (5) | 3 (19) |  |
| 30 to <300 | 3 (14) | 7 (44) |  |
| 300 to <500 | 2 (10) | 2 (12) |  |
| 500 to <1000 | 3 (14) | 1 (6) |  |
| ≥1000 | 12 (57) | 3 (19) |  |
| **Glycemic status** |  |  |  |
| HbA1c category, n (%)‡ | (n=27) | (n=25) | *** |
| <6.5% | 3 (11) | 14 (56) |  |
| 6.5 to <7.5% | 7 (26) | 7 (28) |  |
| 7.5 to <8.5% | 6 (22) | 3 (12) |  |
| ≥8.5% | 11 (41) | 1 (4) |  |
| **Comorbidities & medications** |  |  |  |
| Diabetes history, n (%) | 29 (100) | 17 (63) | *** |
| Diabetes duration, years, median (IQR)† | 22 (16–27) (n=28) | 12 (6–23) (n=16) | * |
| Hypertension history, n (%) | 28 (97) | 26 (96) |  |
| Hypertension duration, years, median (IQR)† | 12 (2–22) (n=27) | 22 (17–30) (n=23) | * |
| On RAAS blockade, n (%) | 29 (100) | 24 (89) |  |

Clinical data obtained from the KPMP Open Access Clinical Data release (March 9, 2026; https://www.kpmp.org/available-data). Age and duration variables in the KPMP open-access dataset are released as 5- or 10-year bins; medians and IQRs were computed using bin midpoints (†). Albuminuria and HbA1c are released as ordinal categorical bins, with proportions calculated among participants with non-missing values (‡). Continuous variables were compared using the Mann-Whitney U test; categorical variables using Fisher's exact test (2x2) or chi-squared test (otherwise). HKD = hypertensive kidney disease; DKD = diabetic kidney disease; eGFR = estimated glomerular filtration rate; RAAS = renin-angiotensin-aldosterone system. \*p<0.05, \*\*p<0.01, \*\*\*p<0.001.

### Proof-of-Concept Analysis in a Small DKD Cohort (n=6)

We first applied our GWR pipeline to a previously published small DKD spatial transcriptomics dataset (n=6; Abedini et al.^7^). All six samples passed sample-level quality control (see STAR Methods). GWR identified spatially heterogeneous MAC-M2-fibrosis coupling in all samples retained after QC, with local regression coefficients varying continuously across tissue space (Figure 2C). High-coupling spots with significantly positive M2 coefficients (BH-adjusted p < 0.05) were observed as compact spatial clusters rather than dispersed spots (Figure 2D; representative sample HK2529, 241 high-coupling spots). High-coupling spots were not restricted to regions with the highest M2 macrophage abundance (Figure 2A) or the highest fibrosis score (Figure 2B). The proportion of high-coupling spots varied substantially across the six samples, consistent with spatial non-stationarity at the cohort level.

**Figure 2.**
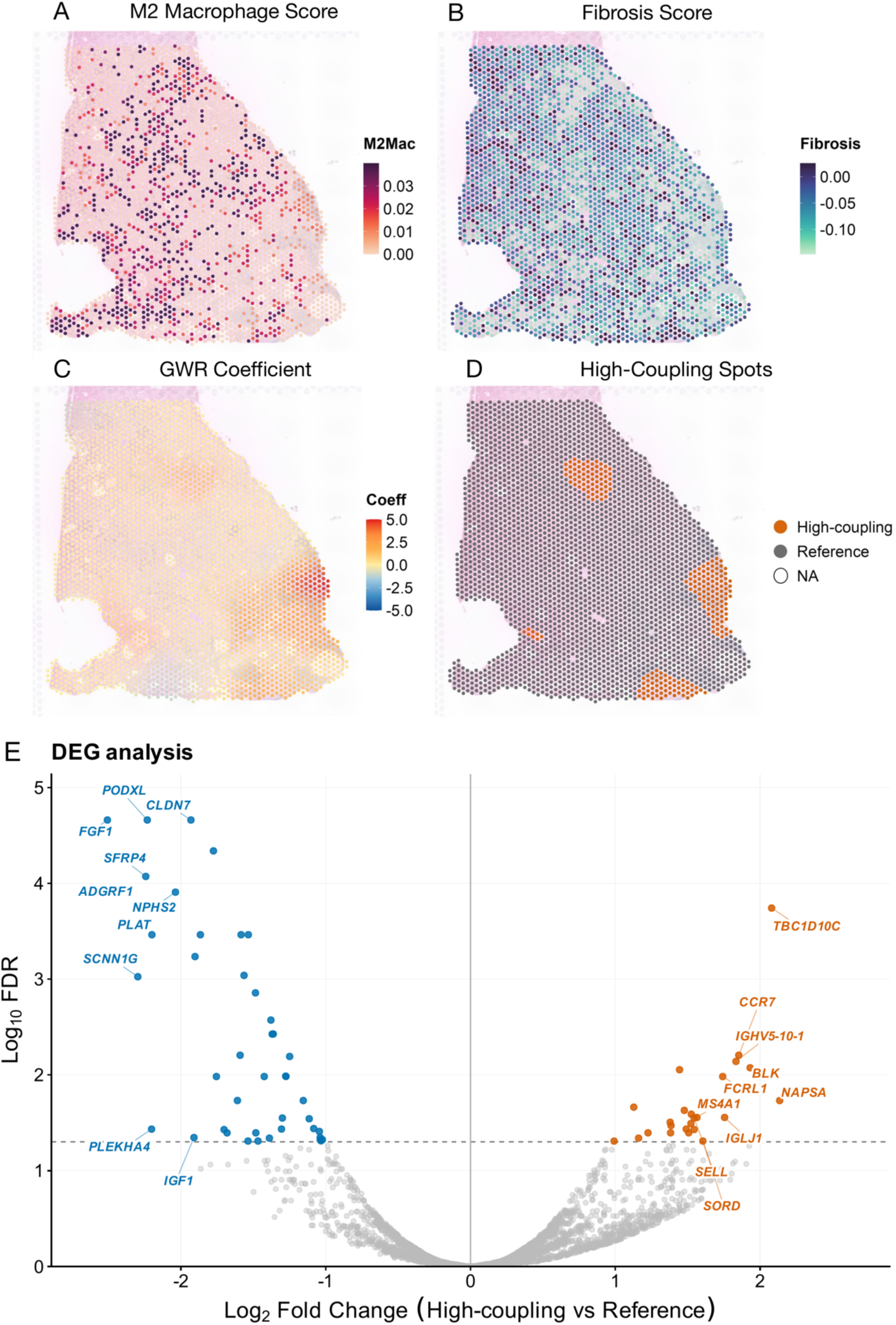
Proof-of-concept validation in a small DKD cohort (n=6^7^) using a representative sample (HK2529). (A) Per-spot M2 macrophage prediction score, estimated by label transfer from the KPMP single-cell RNA-seq reference atlas^8^. Color scale is per-sample; higher values indicate greater predicted M2 macrophage abundance. (B) Per-spot fibrosis score, computed from myofibroblast marker genes using AddModuleScore. Color scale is per-sample. (C) Local GWR regression coefficient (β) for the MAC-M2-fibrosis association at each spot, displayed on a symmetric scale centered at zero. Positive coefficients (orange) indicate local zones where M2 macrophage abundance is positively associated with fibrosis; negative coefficients (blue) indicate inverse local associations. (D) High-coupling spots (vermillion; M2 coefficient > 0 and BH-adjusted p < 0.05) versus reference spots (gray); spots with non-evaluable GWR estimates are shown as open circles (NA). High-coupling regions do not consistently coincide with regions of highest M2 macrophage score (A) or highest fibrosis score (B); rather, they correspond to zones of significant local co-variation between the two variables. (E) Volcano plot of pseudobulk DEGs between GWR-defined high-coupling and reference spots, pooled across the 3 of 6 DKD samples that met the threshold (≥ 20 high-coupling and ≥ 20 reference spots; degrees of freedom = 2). Genes are colored by direction of differential expression: vermillion, upregulated (FDR < 0.05; n = 24); blue, downregulated (FDR < 0.05; n = 43); gray, not significant. Top 10 up- and down-regulated genes by absolute log2 fold-change are labeled. The 67 significant genes (24 up, 43 down) include B cell and plasma cell markers (*BLK*, *BANK1*, *CD79A*, *FCRL1*, *FCMR*, *MS4A1*, *POU2AF1*), immunoglobulin variable- and joining-region genes (*IGHV5-10-1*, *IGHV3-33*, *IGHJ6*, *IGLJ1*), and lymphocyte trafficking factors (*CCR7*, *SELL*) on the upregulated side, consistent with the TLS-like immune aggregates independently described by Abedini et al.^7^ in the DKD fibrotic microenvironment, and three podocyte-specific markers (*NPHS2*, *PODXL*, *CLIC5*), claudin-family tight junction proteins (*CLDN4*, *CLDN7*), and collecting-duct and tubular epithelial markers on the downregulated side, reflecting loss of normal nephron parenchymal identity within high-coupling spots. The complete gene list is provided in Supplementary Table S2.

Of the six DKD samples, three (HK2529, HK2877, HK2844) met the threshold for downstream DEG analysis (at least 20 high-coupling spots and at least 20 reference spots). Spatial maps for the two non-representative DEG-eligible samples (HK2844 and HK2877) are shown in Figure S1. Pseudobulk DEG analysis (edgeR, paired design; degrees of freedom = 2) identified 67 significant genes (FDR < 0.05; 24 upregulated, 43 downregulated; Figure 2E; Supplementary Table S2). Upregulated genes were dominated by B cell and plasma cell markers (*BLK*, *BANK1*, *CD79A*, *FCRL1*, *FCMR*, *MS4A1*, *POU2AF1*, *NIBAN3*, *PARP15*), immunoglobulin variable- and joining-region genes indicative of B cell receptor diversification (*IGHV5-10-1*, *IGHV3-33*, *IGHJ6*, *IGLJ1*), and lymphocyte trafficking factors (*CCR7*, *SELL*), together suggesting local adaptive immune organization. Downregulated genes included three podocyte-specific markers (*NPHS2*, *PODXL*, *CLIC5*), claudin-family tight junction proteins (*CLDN4*, *CLDN7*), collecting-duct and tubular epithelial markers (*AQP2*, *SCNN1A*, *SCNN1G*, *ATP6V0A4*, *CALB1*, *HOPX*), and the anti-inflammatory protease inhibitor *SLPI*, reflecting a broader loss of differentiated nephron parenchymal identity within high-coupling spots.

### Replication: KPMP DKD Cohort (n=30)

To increase statistical power for DEG analysis and to replicate the proof-of-concept findings in an independent cohort, we applied the identical pipeline to the larger KPMP DKD cohort (30 Visium samples; 29 retained after QC; Figure 1C). GWR identified spatially heterogeneous MAC-M2-fibrosis coupling across all samples retained after QC (representative sample 29-10282 shown in Figure 3A–D), with 14 samples meeting the threshold for DEG analysis (at least 20 high-coupling spots and at least 20 reference spots; degrees of freedom = 13). Spatial maps for the remaining 13 DEG-eligible samples are shown in Figures S2 and S3.

**Figure 3.**
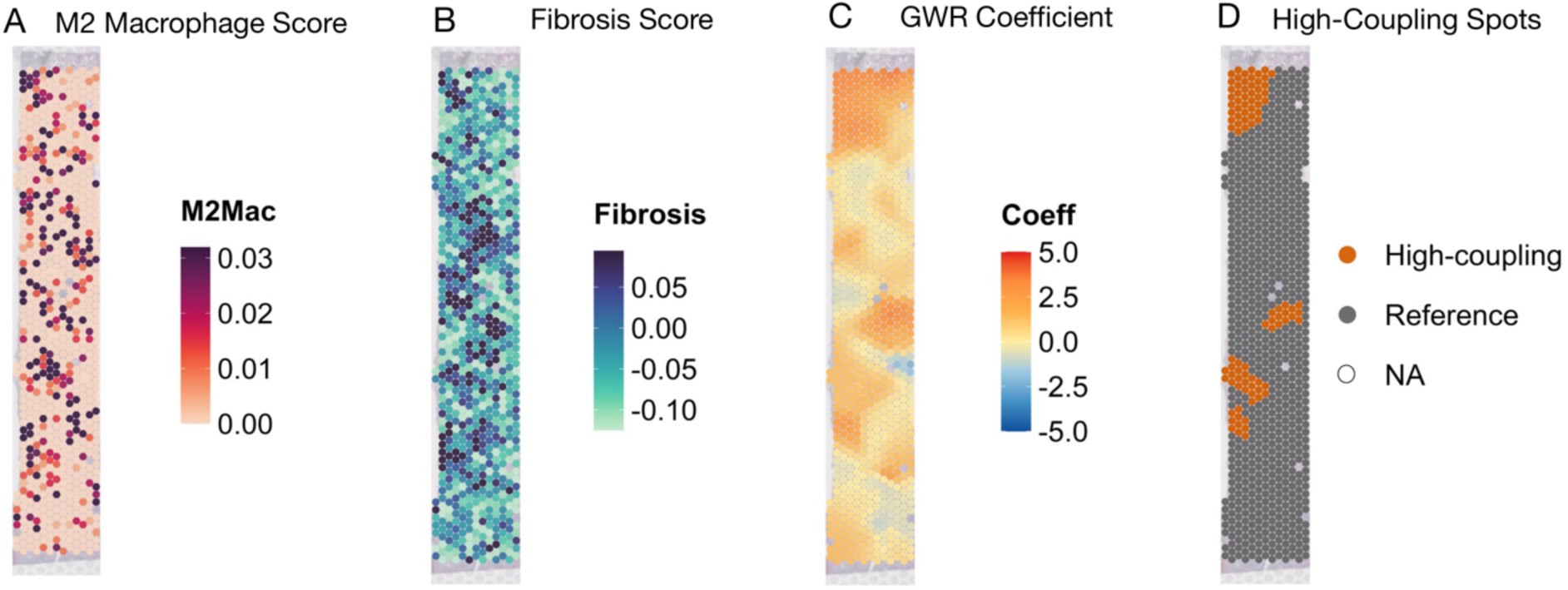
Spatial visualization in a representative KPMP DKD sample (29-10282). (A) Per-spot M2 macrophage prediction score, estimated by label transfer from the KPMP single-cell RNA-seq reference atlas. Color scale is per-sample. (B) Per-spot fibrosis score, computed from myofibroblast marker genes. Color scale is per-sample. (C) Local GWR regression coefficient for the MAC-M2-fibrosis association, displayed on a symmetric scale centered at zero, illustrating spatially varying coupling strength across the tissue. (D) High-coupling spots (vermillion; M2 coefficient > 0 and BH-adjusted p < 0.05) and reference spots (gray); spots with non-evaluable GWR estimates are shown as open circles (NA). High-coupling spots are spatially clustered rather than randomly distributed across the tissue. Note that high-coupling regions in (D) do not consistently coincide with regions of highest M2 macrophage score (A) or highest fibrosis score (B); rather, they correspond to zones of significant local co-variation between the two variables.

DEG analysis identified 128 significant genes (FDR < 0.05; Figure 4A). Genes upregulated in high-coupling spots included IgE-related and mast cell-associated genes, including *IGHE*, *FCER1A*, and the mast cell tryptase *TPSB2*, together with the inflammatory damage-associated molecular pattern (DAMP) *S100A8* (the most significantly upregulated gene) and fibrillar collagens such as *COL1A1*. The most significantly downregulated genes were vascular mural and pericyte markers, led by *DES* (the most statistically significant gene overall), *CNN1*, *ACTG2*, *MYH11*, and *RGS5*, with reduced endothelial markers, indicating loss of mature vascular identity within high-coupling spots. Together, the IgE/mast cell co-enrichment alongside inflammatory and ECM-related signals is consistent with an inflammatory, fibrogenic character of these regions; representative effect sizes are shown in Figure 4A.

**Figure 4.**
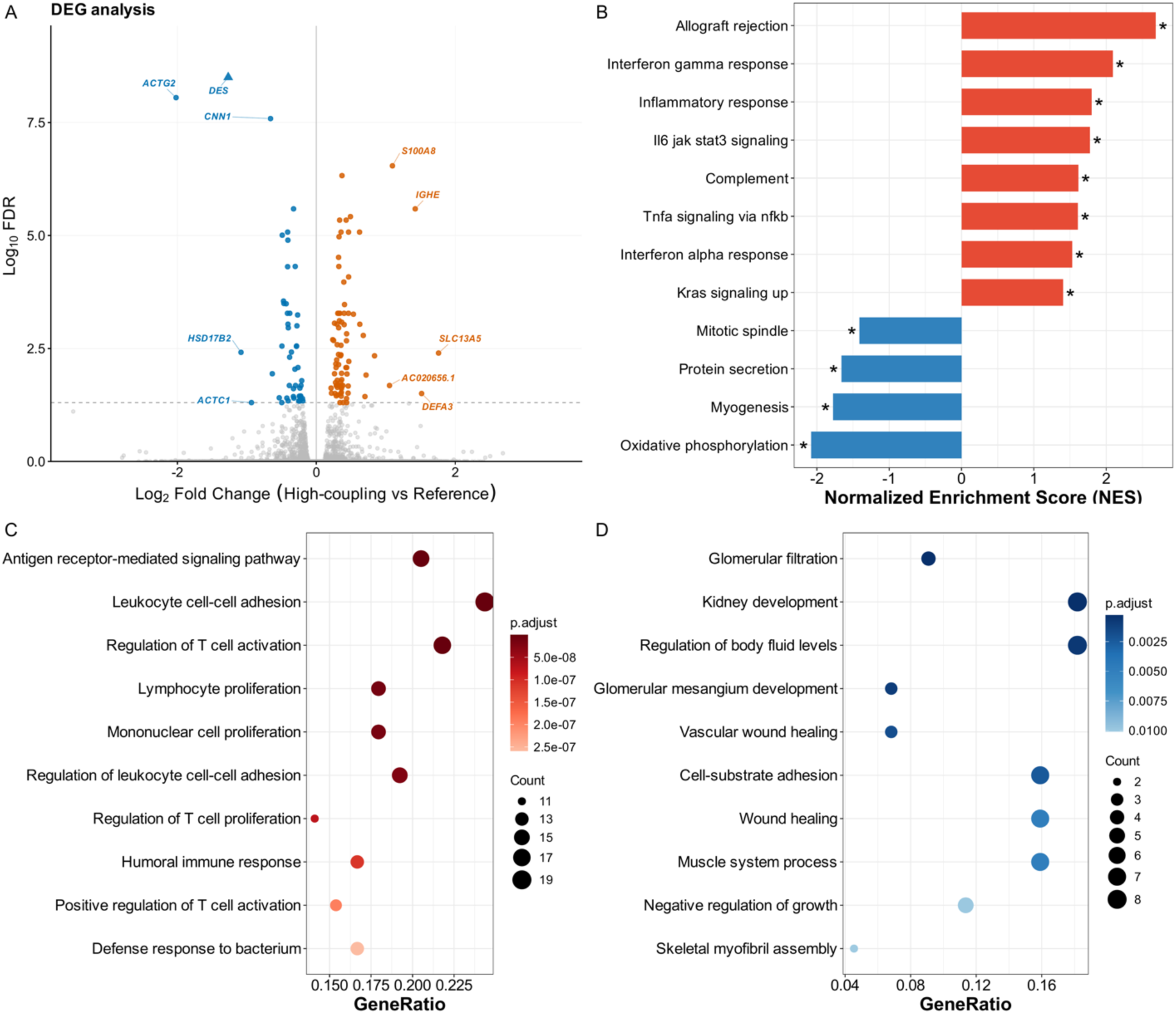
Primary analysis: KPMP DKD cohort differential expression and pathway enrichment (n=30). (A) Volcano plot of pseudobulk DEGs between GWR-defined high-coupling and reference spots. Of 29 QC-passing samples, 14 met the threshold for DEG analysis (degrees of freedom = 13). Genes are colored by direction of differential expression: vermillion, upregulated (FDR < 0.05); blue, downregulated (FDR < 0.05); gray, not significant. Genes meeting FDR < 10⁻⁵ are labeled, prioritizing statistical robustness over fold-change magnitude. *DES* (FDR = 1.5×10⁻¹²) is displayed as a triangle marker at the y-axis cap (y = 9); its actual −log₁₀ FDR is 11.83. (B) GSEA Hallmark pathway analysis ranked by normalized enrichment score (NES). Bar plot shows the top 12 pathways by absolute NES among 13 significantly enriched Hallmark pathways (adjusted p-value < 0.05). Immune and inflammatory pathways are positively enriched, whereas oxidative phosphorylation and myogenesis-related programs are depleted. Asterisks (*) denote adjusted p-value < 0.05. (C) GO biological process enrichment for genes upregulated in high-coupling spots, with redundant terms collapsed using semantic similarity (clusterProfiler::simplify, cutoff = 0.7, by = “p.adjust”). The top 10 non-redundant terms, ordered by adjusted p-value (most significant at top), are shown. Dot size indicates the number of input genes (Count) within each term; color intensity reflects adjusted p-value (darker = more significant). (D) GO biological process enrichment for genes downregulated in high-coupling spots, processed identically to panel C.

GO biological process enrichment (redundant terms collapsed by semantic similarity; STAR Methods) clearly separated the two directions. Upregulated genes were enriched almost exclusively for adaptive immune processes, including antigen receptor-mediated signaling (the top term by adjusted p-value), leukocyte cell-cell adhesion, and regulation of T cell activation (Figure 4C), whereas downregulated genes were enriched for kidney-specific and tissue-remodeling processes, led by glomerular filtration and kidney development (Figure 4D). Consistently, gene set enrichment analysis (GSEA) across Molecular Signatures Database (MSigDB) Hallmark gene sets (13 pathways at adjusted p-value < 0.05; Figure 4B) showed strong positive enrichment of immune and inflammatory programs, with Allograft Rejection the top-ranked pathway (normalized enrichment score (NES) = 2.7), followed by Interferon-Gamma Response, Inflammatory Response, IL6/JAK/STAT3 Signaling, Complement, and TNF-α Signaling via NF-κB, and coordinate depletion of metabolic and contractile programs (Oxidative Phosphorylation, NES = −2.1; Myogenesis, NES = −1.8).

### Disease Specificity: KPMP HKD Cohort (n=27)

In the KPMP HKD cohort, GWR identified spatially heterogeneous MAC-M2-fibrosis coupling (Figure 5A-D), and 9 samples met criteria for DEG analysis (at least 20 high-coupling spots and at least 20 reference spots; degrees of freedom = 8). Spatial maps for the remaining 8 DEG-eligible samples are shown in Figures S4 and S5. DEG analysis revealed 21 significant genes (FDR < 0.05; Table 2). The most strongly upregulated gene was *IGFL1* (logFC = +3.94), followed by *CXCL6*, *SERPINA3*, and *VCAN*. Downregulated genes included collecting duct intercalated cell markers (*AQP6*, *SLC4A1*, *RHCG*, *ATP6V0D2*), smooth muscle markers (*ACTG2*, *MYL9*, *TPM2*, *PLN*), juxtaglomerular renin (*REN*), and immunoglobulin genes (*IGLC7*, *IGHD*). Notably, the IgE-related genes *IGHE* and *FCER1A*, the mast cell tryptase *TPSB2*, and the inflammatory gene *S100A8*, which were significantly upregulated in KPMP DKD high-coupling spots, were not significantly enriched in the HKD signature (all FDR > 0.05), supporting the possibility that these DKD-associated immune genes are more specific to DKD than to kidney fibrosis in general.

**Figure 5.**
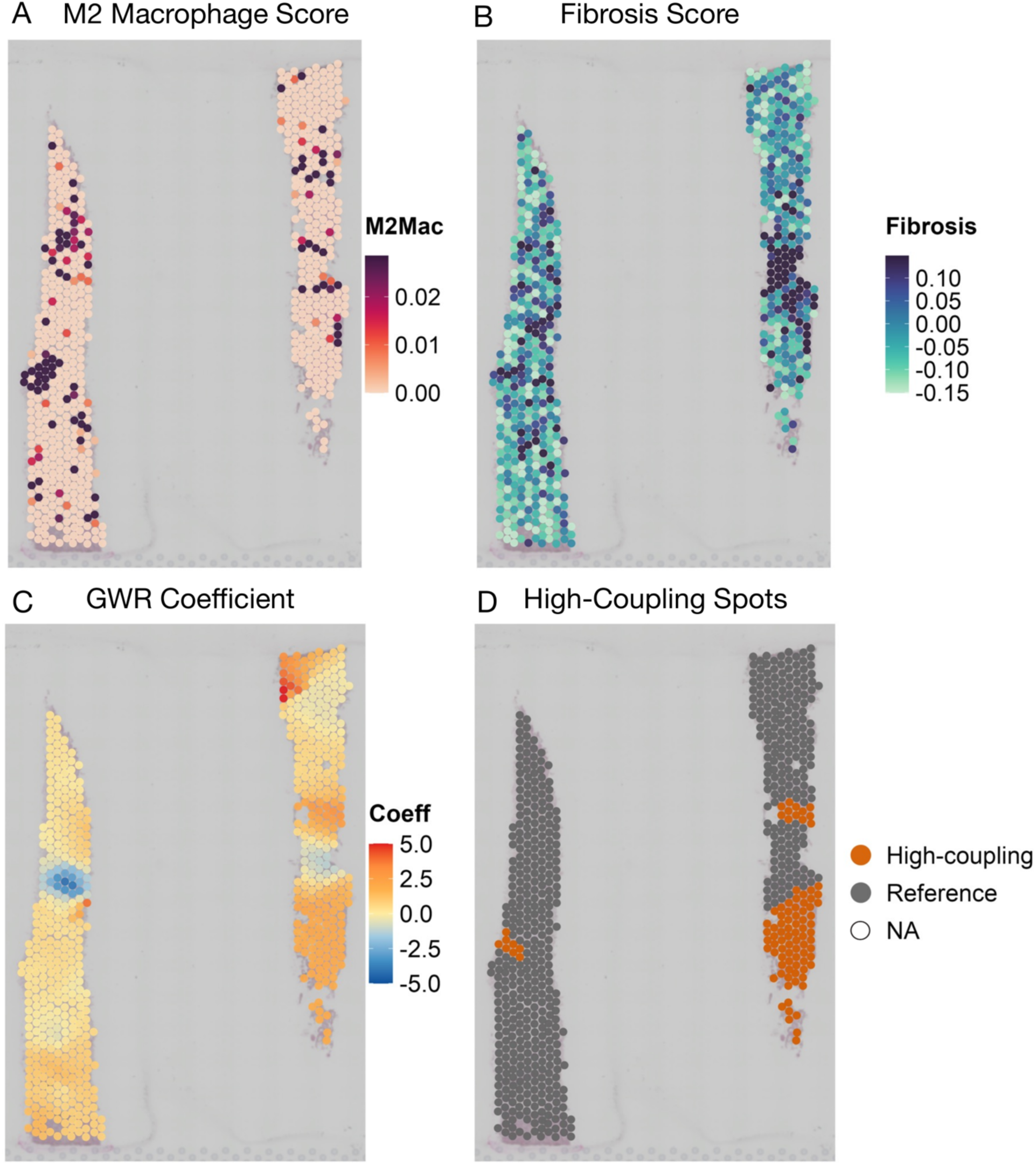
Cross-disease comparison: spatial visualization in the KPMP HKD cohort, shown for a representative sample (28-12518). (A) Per-spot M2 macrophage prediction score. (B) Per-spot fibrosis score. For each continuous score, the color scale was set per sample using the 2.5–97.5 percentile range; values outside this range are shown in the most extreme color. (C) Local GWR regression coefficient for the MAC-M2-fibrosis association, displayed on a per-sample symmetric scale centered at zero. (D) High-coupling spots (vermillion; M2 coefficient > 0 and BH-adjusted p < 0.05) and reference spots (gray); spots with non-evaluable GWR estimates are shown as open circles (NA). As in the KPMP DKD cohort, high-coupling regions in (D) do not consistently coincide with regions of highest M2 macrophage score (A) or highest fibrosis score (B); rather, they correspond to zones of significant local co-variation between the two variables. The full list of significantly differentially expressed genes between high-coupling and reference spots in the KPMP HKD cohort is presented in Table 2.

**Table 2.** Table of all 21 significant genes (FDR < 0.05) identified by pseudobulk DEG analysis between high-coupling and reference spots in the KPMP HKD cohort, with up- and down-regulated genes shown separately. Values are log2 fold-change (HighCoupling/Reference) and Benjamini-Hochberg adjusted FDR. The IgE-related genes *IGHE* and *FCER1A* and the mast cell tryptase *TPSB2*, together with the inflammatory gene *S100A8*, which were significantly upregulated in the KPMP DKD cohort, were not significantly enriched in the HKD signature (all FDR > 0.05).

|  | log<sub>2</sub>FC | FDR |
| --- | --- | --- |
| **Up in high-coupling spots (n = 4)** |  |  |
| *IGFL1* | +3.94 | $3.30 \times 10^{-4}$ |
| *CXCL6* | +0.81 | $1.30 \times 10^{-2}$ |
| *SERPINA3* | +0.65 | $3.88 \times 10^{-2}$ |
| *VCAN* | +0.48 | $3.41 \times 10^{-3}$ |
| **Down in high-coupling spots (n = 17)** |  |  |
| *ACTG2* | -1.66 | $2.52 \times 10^{-5}$ |
| *IGHD* | -1.52 | $1.08 \times 10^{-4}$ |
| *IGLC7* | -1.20 | $5.27 \times 10^{-7}$ |
| *REN* | -1.07 | $7.29 \times 10^{-3}$ |
| *AC019080.1* | -0.84 | $1.19 \times 10^{-5}$ |
| *PLN* | -0.69 | $1.28 \times 10^{-3}$ |
| *SLC39A5* | -0.65 | $3.28 \times 10^{-2}$ |
| *SLC4A1* | -0.62 | $2.36 \times 10^{-5}$ |
| *DMRT2* | -0.62 | $8.47 \times 10^{-3}$ |
| *AQP6* | -0.54 | $8.47 \times 10^{-3}$ |
| *RHCG* | -0.52 | $3.08 \times 10^{-3}$ |
| *ATP6V0D2* | -0.51 | $9.70 \times 10^{-3}$ |
| *ADIRF* | -0.50 | $2.36 \times 10^{-5}$ |
| *IGFBP3* | -0.50 | $1.25 \times 10^{-4}$ |
| *TPM2* | -0.45 | $3.62 \times 10^{-3}$ |
| *ECRG4* | -0.41 | $3.10 \times 10^{-2}$ |
| *MYL9* | -0.37 | $3.28 \times 10^{-2}$ |

### Cross-cohort observations from spatial visualizations

Visual inspection of spatial maps across all three cohorts (Figures 2A-D, 3A-D, 5A-D, and Figures S1-S5) revealed two recurring patterns. First, in samples that yielded sufficient high-coupling spots for downstream DEG analysis, high-coupling spots formed discrete tissue clusters rather than being randomly distributed, indicating that GWR identifies spatially organized microenvironments rather than stochastic noise. Second, these high-coupling regions did not consistently coincide with areas of highest M2 macrophage score or highest fibrosis score in isolation; instead, they corresponded to zones where M2 abundance and fibrosis score co-varied at the local scale, including regions where both scores were intermediate but tightly linked. This pattern is consistent with the conceptual basis of GWR, which detects local co-variation rather than absolute co-elevation of two variables.

## Discussion

We applied GWR to kidney spatial transcriptomics and identified spatially restricted regions in DKD where M2 macrophage abundance was locally associated with fibrosis. These GWR-defined high-coupling spots were not simply areas with high M2 macrophage abundance or high fibrosis score alone. Rather, they marked regions in which the two signals varied together at the local scale. The main findings are that GWR can capture spatially varying MAC-M2-fibrosis associations in human kidney tissue, that DKD high-coupling regions show adaptive immune activation with IgE-related immune features, and that these features are not observed in the same way in HKD.

A major motivation for this study was the limitation of conventional spatial transcriptomic analyses in addressing local relationships between biological variables. Many current approaches map cell-type localization, test co-localization, or classify tissue microenvironments. These analyses are useful for describing tissue organization, but they do not directly quantify whether the relationship between two continuous variables varies across tissue space. GWR addresses this question by estimating a local regression coefficient for each spot. In our analysis, this coefficient represented the local association between M2 macrophage abundance and fibrosis score, after adjustment for technical variation in sequencing depth. This differs from global regression, which assumes a single tissue-wide association, and from co-localization analysis, which asks whether two signals occupy the same area. By assigning a local coefficient to each spot, GWR allowed us to define regions where the MAC-M2-fibrosis association was significantly positive.

One informative feature of the spatial maps was that high-coupling regions did not consistently coincide with the regions of highest M2 macrophage score or highest fibrosis score. This distinction is important for interpreting what GWR detects. Areas of severe fibrosis may represent relatively end-stage scarring, where active remodeling has decreased and local variation in fibrosis is limited. In contrast, high-coupling regions may represent tissue areas in which M2 macrophage abundance and fibrosis are still changing together. These regions can include spots with intermediate levels of both signals, as long as their local association is strong. This pattern would not be captured by simple thresholding of M2 macrophage abundance or fibrosis score. The clustered distribution of high-coupling spots further supports the idea that they correspond to spatially organized microenvironments rather than random spot-level noise.

Only a subset of samples contained enough high-coupling and reference spots for downstream DEG analysis: 3 of 6 samples in the proof-of-concept DKD cohort, 14 of 29 samples in the KPMP DKD cohort, and 9 of 23 samples in the KPMP HKD cohort. We do not view the remaining samples simply as technical failures. Instead, this pattern likely reflects differences in tissue composition, disease stage, and the extent to which M2 macrophages and fibrosis are locally coupled within each biopsy. Samples with homogeneous or advanced fibrosis may have limited spatial variation in fibrosis score, reducing the ability of GWR to identify discrete regions of positive local association.

Biopsies containing both preserved and injured parenchyma may be more likely to contain detectable high-coupling regions. Thus, GWR-defined high-coupling spots may preferentially capture spatially restricted remodeling microenvironments rather than uniformly scarred tissue.

The molecular features of DKD high-coupling spots pointed to adaptive immune activation in both DKD cohorts. In the proof-of-concept cohort, high-coupling spots were enriched for B cell and plasma cell markers, including *BLK*, *BANK1*, *CD79A*, *FCRL1*, *FCMR*, *MS4A1*, and *POU2AF1*, together with immunoglobulin variable- and joining-region genes such as *IGHV5-10-1*, *IGHV3-33*, *IGHJ6*, and *IGLJ1*. Lymphocyte trafficking genes, including *CCR7* and *SELL*, were also upregulated. These findings are consistent with the tertiary lymphoid structure-like immune aggregates described by Abedini et al.^7^ in the DKD fibrotic microenvironment and support the biological relevance of the GWR-defined regions. Downregulated genes in this cohort included podocyte markers such as *NPHS2*, *PODXL*, and *CLIC5*, raising the possibility that glomerular injury may be spatially linked to regions of strong MAC-M2-fibrosis coupling.

In the larger KPMP DKD cohort, high-coupling spots showed a related but more specific immune signature. Upregulated genes included *IGHE*, *FCER1A*, and the mast cell tryptase *TPSB2*, together with the inflammatory DAMP *S100A8* and the B cell-organizing chemokine *CCL19*. Pathway analyses supported this immune-activated state. GO biological process terms enriched among upregulated genes included antigen receptor-mediated signaling, leukocyte cell-cell adhesion, and regulation of T cell activation. Hallmark GSEA identified Allograft Rejection as the top-ranked pathway, along with Interferon-Gamma Response, Inflammatory Response, and IL6/JAK/STAT3 Signaling. These results suggest that high-coupling spots in DKD are not only fibrotic regions, but immune-active fibrotic microenvironments.

The IgE-related signal in the KPMP DKD cohort is consistent with prior work implicating mast cells and FcεRI-related pathways in kidney fibrosis and DKD^10^. Mast cells are rare in normal kidney cortex but increase in fibrotic kidney disease, including DKD, and their abundance has been reported to correlate with interstitial fibrosis across nephropathies^11,12^. Upon IgE-FcεRI activation, mast cells can release mediators such as TGF-β, chymase, tryptase, TNF-α, renin, and histamine, which may promote fibroblast activation, extracellular matrix deposition, and myofibroblast differentiation^13,14^. Prior bulk transcriptomic analysis also reported increased *FCER1A* expression in human DKD glomeruli and proposed the IgE receptor as a potential therapeutic target in DKD^15^. Because those studies used bulk or non-spatial approaches, they could not determine where these signals arise within kidney tissue. Our results add spatial context by showing that *IGHE*, *FCER1A*, and *TPSB2* are enriched within GWR-defined high-coupling spots, suggesting that IgE-related immune responses may be a feature of DKD fibrotic microenvironments in which M2 macrophages are locally associated with fibrosis.

One possible model is that M2 macrophages and IgE-related immune activity reinforce each other within local fibrotic niches. M2 macrophage-associated cytokines such as IL-4 and IL-13 could support IgE-related immune responses, while mast cell-derived mediators such as TGF-β and tryptase could promote extracellular matrix remodeling and further macrophage polarization. This model remains speculative, and the present data do not establish a causal circuit. However, the spatial restriction of *IGHE*, *FCER1A*, and *TPSB2* to high-coupling regions provides a rationale for testing whether IgE-related immune responses contribute to DKD fibrosis in future experimental studies.

The two DKD cohorts differed in the specific genes that were most prominent, but both pointed to adaptive immune organization within regions of MAC-M2-fibrosis coupling. In the proof-of-concept cohort, the signal was dominated by B cell and TLS-like markers. In the KPMP cohort, it included IgE-related genes and FcεRI-associated components, together with *CCL19*. These differences may reflect statistical power, cohort composition, tissue sampling, or technical differences between datasets. Nevertheless, the convergence on adaptive immune activation across independent DKD cohorts supports the interpretation that GWR-defined high-coupling spots identify biologically meaningful immune-fibrotic microenvironments. This convergence is further supported by a recent single-cell and spatial atlas of DKD that independently identified a B cell-rich patient subgroup associated with disease progression^16^, consistent with the adaptive immune and B cell/TLS-like signatures enriched within our GWR-defined high-coupling regions.

Another consistent feature across DKD cohorts was the depletion of vascular smooth muscle and pericyte markers, including *DES*, *CNN1*, *ACTG2*, *MYH11*, and *RGS5*. In the KPMP cohort, this pattern extended to additional pericyte and endothelial markers such as *MCAM*, *CSPG4*, *EMCN*, and *CD34*, and was accompanied by upregulation of remodeling-associated matrix genes including *VCAN*, *MMP9*, and *THBS2*. This combination suggests vascular destabilization and extracellular matrix remodeling within high-coupling regions. Given that pericyte loss can drive peritubular capillary rarefaction and tubular injury^17^, and that peritubular capillary rarefaction contributes to CKD progression^18^, vascular injury may be closely linked to the immune-fibrotic state captured by GWR.

The comparison with HKD suggested that the IgE-related immune signature is not simply a general feature of fibrotic kidney tissue. Applying the same pipeline to the KPMP HKD cohort identified high-coupling spots, but their transcriptomic features differed from those in DKD. HKD high-coupling spots showed depletion of collecting duct intercalated cell markers, including *AQP6*, *SLC4A1*, and *RHCG*, as well as smooth muscle genes. In contrast, the DKD IgE-related genes, including *IGHE*, *FCER1A*, and *TPSB2*, were not enriched. These findings suggest that DKD and HKD may share the downstream feature of local MAC-M2-fibrosis coupling while differing in the upstream niche biology associated with that coupling. The loss of collecting duct intercalated cell markers in HKD may reflect disease-specific tubular or acid-base regulatory injury in hypertensive nephrosclerosis, although this interpretation will require further study.

More broadly, this study illustrates how spatial statistics can be used to ask questions that are difficult to address with conventional spatial transcriptomic workflows. In many diseases, the key question is not only whether two signals co-localize, but where their relationship is locally strengthened. Tumor microenvironments, fibrotic organs, and chronically inflamed tissues all contain spatially restricted interactions that may be missed by global or purely descriptive analyses. GWR provides one approach for mapping such local relationships between continuous biological variables. Future studies combining spatial transcriptomics with spatial proteomics, such as MERFISH or Xenium, will be needed to validate IgE and FcεRI-related signals at the protein level and to define the spatial relationships among mast cells, M2 macrophages, and myofibroblasts at single-cell resolution.

### Limitations of the Study

This study has several limitations. First, GWR assumes a local linear relationship between M2 macrophage abundance and fibrosis score. This may not capture non-linear or threshold-dependent interactions, and the local estimates depend on bandwidth selection. Second, Visium data are measured at 55-μm spot resolution, and each spot can contain multiple cell types. High-coupling spots should therefore be interpreted as multicellular microenvironments rather than single-cell states, and the local MAC-M2–fibrosis coupling identified by GWR reflects local co-variation between inferred M2 macrophage abundance and the fibrosis score rather than a direct cellular or lineage relationship. Third, M2 macrophage abundance was inferred by deconvolution, and these estimates depend on the reference atlas and the assumptions of label transfer. Fourth, the IgE-related immune genes identified here, including *TPSB2*, were detected at the transcriptomic level. Protein-level validation, for example by co-immunofluorescence staining for IgE, tryptase, and fibrosis markers such as collagen or α-SMA, will be needed to confirm their localization within GWR-defined high-coupling regions. Finally, the DKD-HKD comparison was cross-sectional and observational. The observed differences between DKD and HKD high-coupling spots may reflect disease-specific mechanisms, differences in disease stage, or other clinical differences. Longitudinal datasets and experimental validation will be required to clarify causality.

## Supporting information

Figure S1. GWR-based spatial analysis in Proof-of-concept DKD samples

Figure S2. GWR-based spatial analysis in KPMP DKD samples (part 1)

Figure S3. GWR-based spatial analysis in KPMP DKD samples (part 2)

Figure S4. GWR-based spatial analysis in KPMP HKD samples (part 1)

Figure S5. GWR-based spatial analysis in KPMP HKD samples (part 2)

Supplementary Table S1. Per-sample QC summary

Supplementary Table S2. Proof-of-concept DKD significant genes

Supplementary Table S3. KPMP DKD significant genes

Supplementary Table S4. GO BP enrichment (upregulated)

Supplementary Table S5. GO BP enrichment (downregulated)

## Resource Availability

### Data and Code Availability

KPMP data: available at https://atlas.kpmp.org/.

Dr. Katalin Susztak Lab data: Abedini et al.^7^; available at http://susztaklab.com and in the Gene Expression Omnibus under accession GSE211785. Analysis code will be made available before publication.

## Acknowledgments

We thank the KPMP consortium, including Dr. Mathias Kretzler, and the Dr. Katalin Susztak Lab for making spatial transcriptomics data available.

## Author Contributions

K.T., H.K. and I.M. conceived the study. K.T. performed the analyses, interpreted the data, and wrote the manuscript. H.K. provided guidance on the analytical strategy and data interpretation. M.N. supervised the study and provided overall scientific advice. I.M. supervised the study and contributed to manuscript writing and revision. All authors reviewed and approved the final manuscript.

## STAR Methods

### Key Resources Table

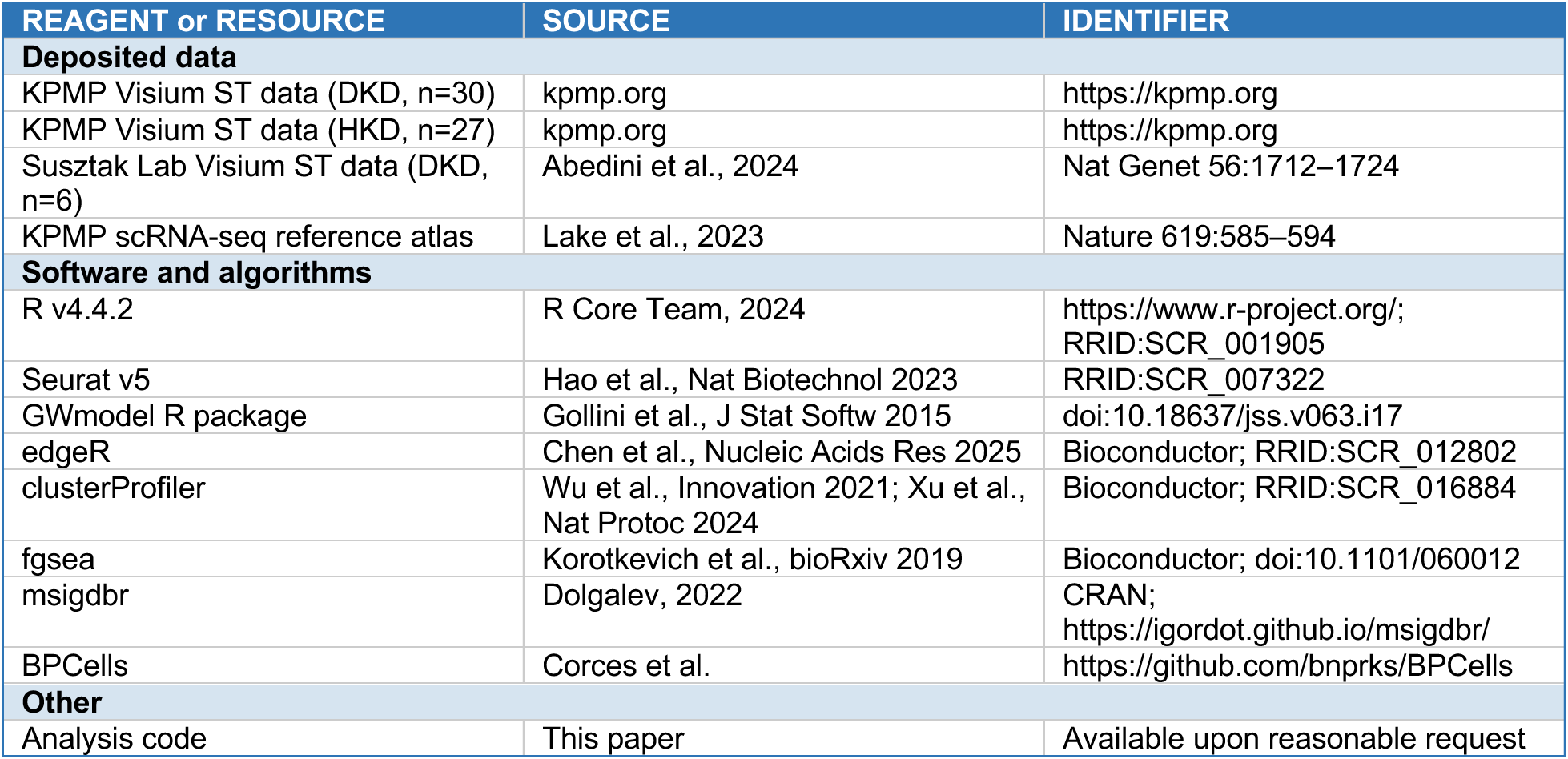

### Method Details

Spatial transcriptomics data processing: Three Visium spatial transcriptomics cohorts were analyzed. The primary cohort comprised 30 KPMP DKD samples (29 retained after QC). The cross-disease cohort comprised 27 KPMP HKD samples (23 retained after QC). The independent validation cohort comprised 6 Susztak Lab DKD samples (all 6 retained after QC). KPMP data were obtained from the open-access KPMP repository (kpmp.org). Susztak Lab data were obtained from Abedini et al.^7^. Raw 10x Visium count matrices were loaded using Load10X_Spatial in Seurat v5^19^. QC was performed at two levels. At the sample level, samples with median nFeature_Spatial < 500, median mitochondrial percentage > 25%, or fewer than 100 spots were excluded. At the spot level, spots with nFeature_Spatial ≤ 500 or mitochondrial percentage ≥ 25% were removed. A per-sample summary of quality-control metrics is provided in Supplementary Table S1.

Normalization and dimensionality reduction: Gene expression counts were normalized using LogNormalize (scale factor = 10,000). After merging samples with JoinLayers, the top 2,000 highly variable genes were identified (vst method), data were scaled, and principal component analysis (PCA) was performed (assay = “Spatial”). Sample names were standardized using make.names().

Cell type deconvolution: Cell type composition was estimated using label transfer with the KPMP single-cell RNA-seq reference atlas^8^. The reference was processed with LogNormalize, top 3,000 highly variable genes, scaling, and PCA. Anchors were identified using FindTransferAnchors (normalization.method = “LogNormalize”, dims = 1:30), and cell type scores were transferred using TransferData with weight.reduction = “pca”. M2 macrophage fraction per spot was defined as the prediction score for the MAC.M2 subclass.

Fibrosis score: A fibrosis score was computed per spot using AddModuleScore. The gene set was derived from the KPMP single-cell reference atlas by identifying myofibroblast (MYOF) marker genes using FindMarkers (Wilcoxon rank-sum test, logfc.threshold = 0.5, min.pct = 0.25, p_val_adj < 0.05), retaining the top genes by average log2FC. The same gene set was applied consistently across all three cohorts. This myofibroblast-derived fibrosis score was used as a spatial readout of fibrotic remodeling, not to infer the cellular origin of myofibroblasts.

Geographically weighted regression: GWR was applied independently to each sample using the GWmodel R package^20^. The model was: Fibrosis_Score ∼ M2_Mac_fraction + log10(unique molecular identifier (UMI) + 1). An adaptive bandwidth was selected by Akaike information criterion (AIC) minimization (bw.gwr) using a bisquare spatial weighting kernel, and spot-level coefficients were estimated with gwr.basic() under the same bisquare kernel. Samples with fewer than 50 spots were excluded. For each spot, a t-value was extracted for the M2 coefficient, and a two-sided p-value was computed from the normal distribution. P-values were adjusted within each sample using the Benjamini-Hochberg method. Spots with M2 coefficient > 0 and BH-adjusted p < 0.05 were classified as high-coupling spots; all remaining spots were classified as reference spots.

Differential gene expression analysis: Samples with at least 20 high-coupling spots and at least 20 reference spots were included in DEG analysis. Pseudobulk counts were aggregated per sample per condition (HighCoupling/Reference) using AggregateExpression (assay = “Spatial”, slot = “counts”). DEG analysis was performed using edgeR^21^ with a paired design: design = ∼ SampleID + Condition. Dispersion was estimated with estimateDisp (robust = TRUE) and quasi-likelihood F-tests were performed with glmQLFTest. Genes with FDR < 0.05 were considered significant.

Enrichment analysis: Gene Ontology Biological Process (GO BP) enrichment was performed using enrichGO (clusterProfiler^22^), with ENTREZID mapping via bitr (org.Hs.eg.db). The universe was restricted to all genes tested in the edgeR differential expression analysis to avoid inflation of enrichment statistics by genes absent from the dataset. Enrichment was performed separately for upregulated and downregulated significant genes. Redundant GO terms were collapsed by semantic similarity using clusterProfiler::simplify (cutoff = 0.7, by = “p.adjust”, select_fun = min), and the top 10 non-redundant terms per direction, ordered by adjusted p-value, were displayed in the main figures. Full pre-simplification and post-simplification enrichment tables are provided in Supplementary Tables S4 (upregulated) and S5 (downregulated). GSEA was performed using fgsea^23^ with Hallmark gene sets (MSigDB, msigdbr). Genes were ranked by sign(logFC) × √F, with a small random jitter added for tie-breaking (set.seed(42)). Pathways with an adjusted p-value < 0.05 were considered significant (minSize = 15, maxSize = 500).

### Quantification and Statistical Analysis

Multiple testing: Benjamini-Hochberg. GWR bandwidth: AIC optimization. All analyses in R 4.4.2

## References

1. Alicic, R.Z., Rooney, M.T., and Tuttle, K.R. (2017). Diabetic Kidney Disease: Challenges, Progress, and Possibilities. Clin J Am Soc Nephrol 12, 2032–2045. 10.2215/cjn.11491116.

2. Humphreys, B.D. (2018). Mechanisms of Renal Fibrosis. Annu Rev Physiol 80, 309–326. 10.1146/annurev-physiol-022516-034227.

3. Schumacher, A., Rookmaaker, M.B., Joles, J.A., Kramann, R., Nguyen, T.Q., van Griensven, M., and LaPointe, V.L.S. (2021). Defining the variety of cell types in developing and adult human kidneys by single-cell RNA sequencing. NPJ Regen Med 6, 45. 10.1038/s41536-021-00156-w.

4. Kuppe, C., Ibrahim, M.M., Kranz, J., Zhang, X., Ziegler, S., Perales-Paton, J., Jansen, J., Reimer, K.C., Smith, J.R., Dobie, R., et al. (2021). Decoding myofibroblast origins in human kidney fibrosis. Nature 589, 281–286. 10.1038/s41586-020-2941-1.

5. Wynn, T.A., and Vannella, K.M. (2016). Macrophages in Tissue Repair, Regeneration, and Fibrosis. Immunity 44, 450–462. 10.1016/j.immuni.2016.02.015.

6. Tang, P.M., Nikolic-Paterson, D.J., and Lan, H.Y. (2019). Macrophages: versatile players in renal inflammation and fibrosis. Nat Rev Nephrol 15, 144–158. 10.1038/s41581-019-0110-2.

7. Abedini, A., Levinsohn, J., Klötzer, K.A., Dumoulin, B., Ma, Z., Frederick, J., Dhillon, P., Balzer, M.S., Shrestha, R., Liu, H., et al. (2024). Single-cell multi-omic and spatial profiling of human kidneys implicates the fibrotic microenvironment in kidney disease progression. Nat Genet 56, 1712–1724. 10.1038/s41588-024-01802-x.

8. Lake, B.B., Menon, R., Winfree, S., Hu, Q., Melo Ferreira, R., Kalhor, K., Barwinska, D., Otto, E.A., Ferkowicz, M., Diep, D., et al. (2023). An atlas of healthy and injured cell states and niches in the human kidney. Nature 619, 585–594. 10.1038/s41586-023-05769-3.

9. Zormpas, E., Queen, R., Comber, A., and Cockell, S.J. (2023). Mapping the transcriptome: Realizing the full potential of spatial data analysis. Cell 186, 5677–5689. 10.1016/j.cell.2023.11.003.

10. Peng, Q.Y., An, Y., Jiang, Z.Z., and Xu, Y. (2024). The Role of Immune Cells in DKD: Mechanisms and Targeted Therapies. J Inflamm Res 17, 2103–2118. 10.2147/jir.S457526.

11. Roberts, I.S., and Brenchley, P.E. (2000). Mast cells: the forgotten cells of renal fibrosis. J Clin Pathol 53, 858–862. 10.1136/jcp.53.11.858.

12. Zheng, J.M., Yao, G.H., Cheng, Z., Wang, R., and Liu, Z.H. (2012). Pathogenic role of mast cells in the development of diabetic nephropathy: a study of patients at different stages of the disease. Diabetologia 55, 801–811. 10.1007/s00125-011-2391-2.

13. Kondo, S., Kagami, S., Kido, H., Strutz, F., Müller, G.A., and Kuroda, Y. (2001). Role of mast cell tryptase in renal interstitial fibrosis. J Am Soc Nephrol 12, 1668–1676. 10.1681/asn.V1281668.

14. Owens, E.P., Vesey, D.A., Kassianos, A.J., Healy, H., Hoy, W.E., and Gobe, G.C. (2019). Biomarkers and the role of mast cells as facilitators of inflammation and fibrosis in chronic kidney disease. Transl Androl Urol 8, S175–s183. 10.21037/tau.2018.11.03.

15. Sur, S., Nguyen, M., Boada, P., Sigdel, T.K., Sollinger, H., and Sarwal, M.M. (2021). FcER1: A Novel Molecule Implicated in the Progression of Human Diabetic Kidney Disease. Front Immunol 12, 769972. 10.3389/fimmu.2021.769972.

16. Dumoulin, B., Levinsohn, J., Klötzer, K.A., Li, C., Mao, L., Ha, E., Mohandes, S., Nguyen, T., Paruzzo, L., Hirohama, D., et al. (2026). Spatial atlas of diabetic kidney disease reveals a B cell-rich subgroup. Nature 653, 1158–1169. 10.1038/s41586-026-10363-4.

17. Kramann, R., Wongboonsin, J., Chang-Panesso, M., Machado, F.G., and Humphreys, B.D. (2017). Gli1(+) Pericyte Loss Induces Capillary Rarefaction and Proximal Tubular Injury. J Am Soc Nephrol 28, 776–784. 10.1681/asn.2016030297.

18. Kida, Y. (2020). Peritubular Capillary Rarefaction: An Underappreciated Regulator of CKD Progression. Int J Mol Sci 21. 10.3390/ijms21218255.

19. Hao, Y., Stuart, T., Kowalski, M.H., Choudhary, S., Hoffman, P., Hartman, A., Srivastava, A., Molla, G., Madad, S., Fernandez-Granda, C., and Satija, R. (2024). Dictionary learning for integrative, multimodal and scalable single-cell analysis. Nat Biotechnol 42, 293–304. 10.1038/s41587-023-01767-y.

20. Gollini, I., Lu, B., Charlton, M., Brunsdon, C., and Harris, P. (2015). GWmodel: An R Package for Exploring Spatial Heterogeneity Using Geographically Weighted Models. Journal of Statistical Software 63, 1–50. 10.18637/jss.v063.i17.

21. Chen, Y., Chen, L., Lun, A.T.L., Baldoni, P.L., and Smyth, G.K. (2025). edgeR v4: powerful differential analysis of sequencing data with expanded functionality and improved support for small counts and larger datasets. Nucleic Acids Res 53. 10.1093/nar/gkaf018.

22. Wu, T., Hu, E., Xu, S., Chen, M., Guo, P., Dai, Z., Feng, T., Zhou, L., Tang, W., Zhan, L., et al. (2021). clusterProfiler 4.0: A universal enrichment tool for interpreting omics data. Innovation (Camb) 2, 100141. 10.1016/j.xinn.2021.100141.

23. Korotkevich, G., Sukhov, V., Budin, N., Shpak, B., Artyomov, M.N., and Sergushichev, A. (2019). Fast gene set enrichment analysis. bioRxiv. 10.1101/060012.

