## Supplementary figures and images for "Applying Spatial Statistics to Spatial Transcriptomics Reveals Local Association Between M2-like Macrophages and Fibrosis in Diabetic Kidney Disease"

### Figure S1. GWR-based spatial analysis in Proof-of-concept DKD samples

A

HK2844

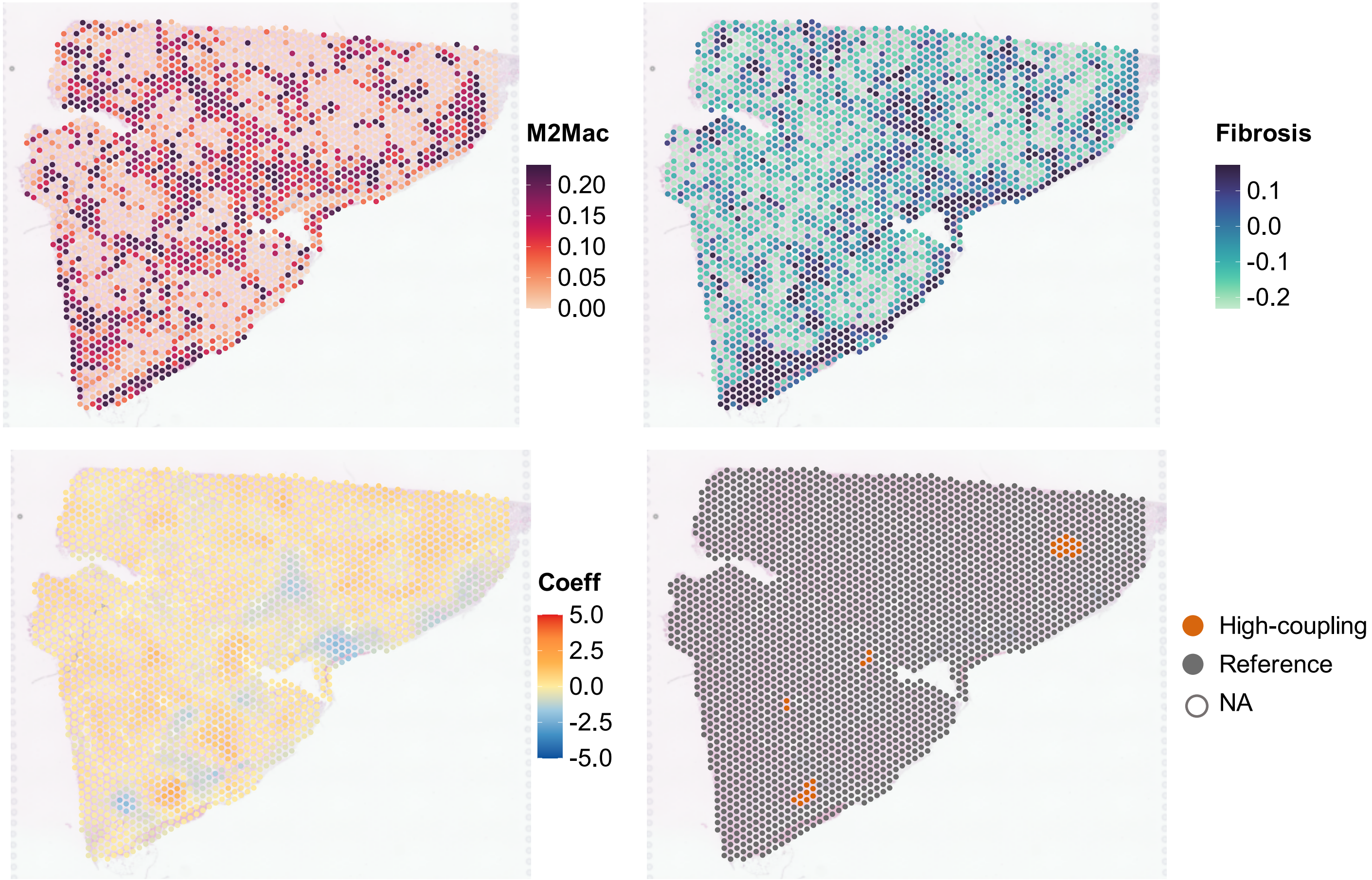


B

HK2877

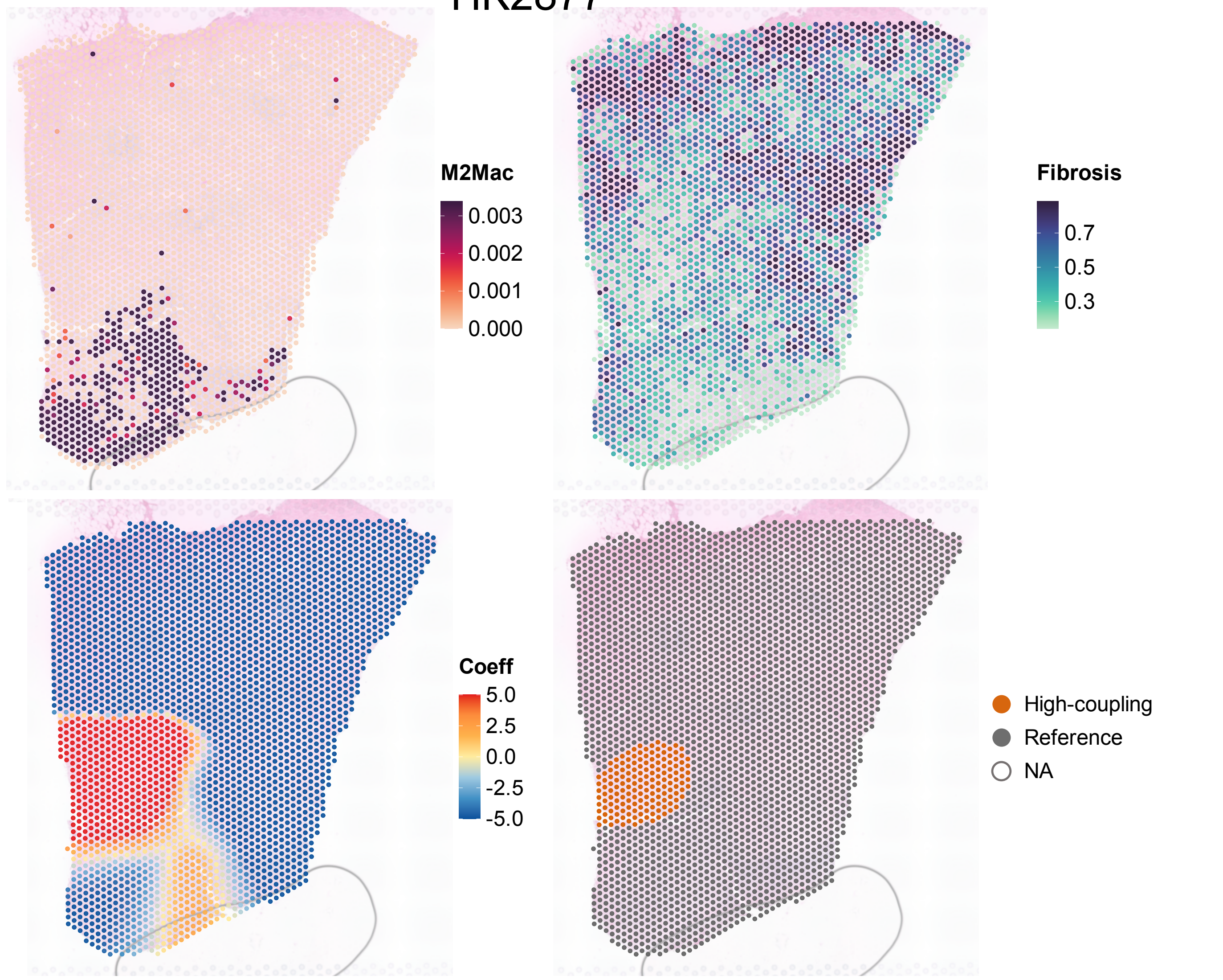

### Figure S2. GWR-based spatial analysis in KPMP DKD samples (part 1)

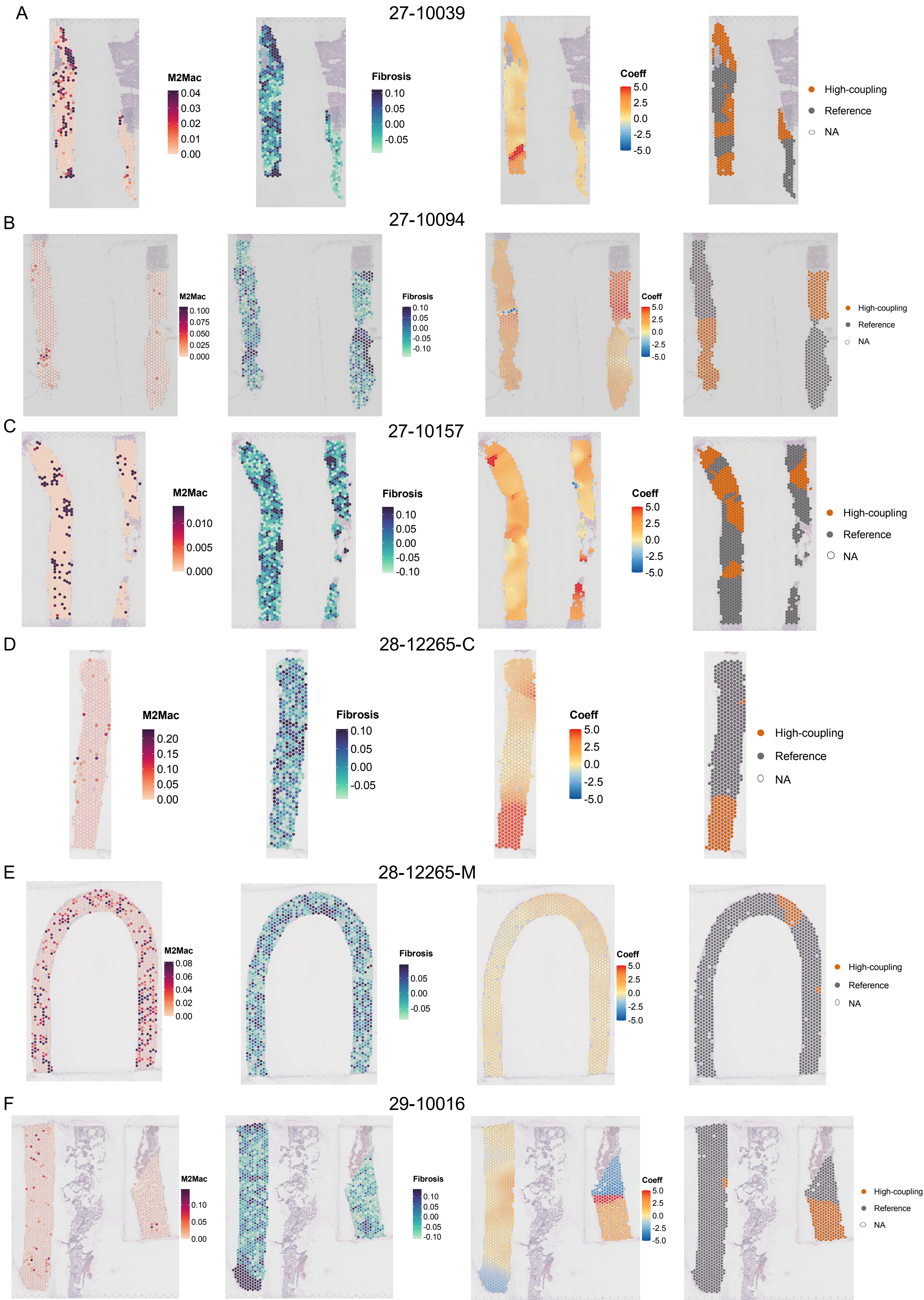

### Figure S3. GWR-based spatial analysis in KPMP DKD samples (part 2)

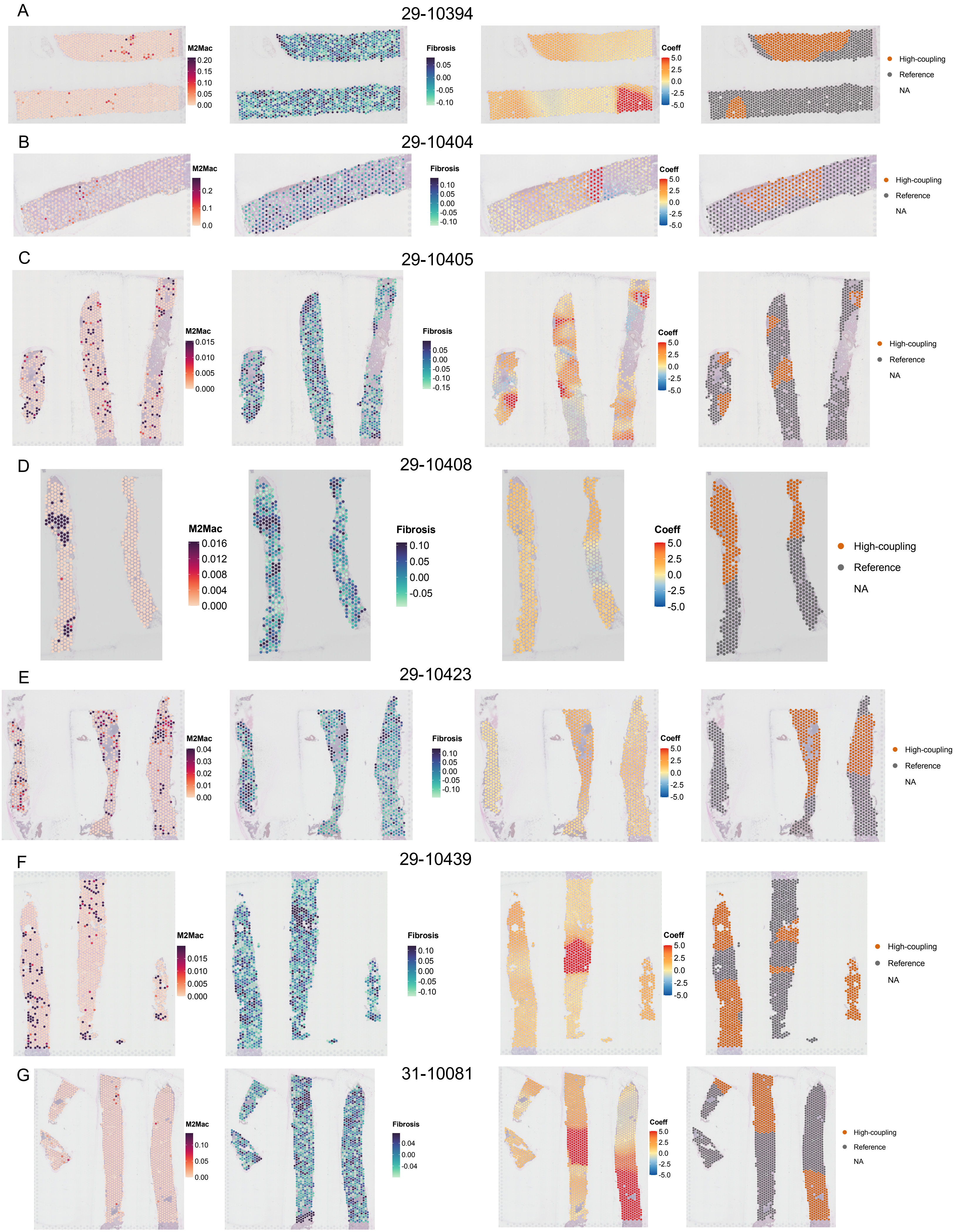

### Figure S4. GWR-based spatial analysis in KPMP HKD samples (part 1)

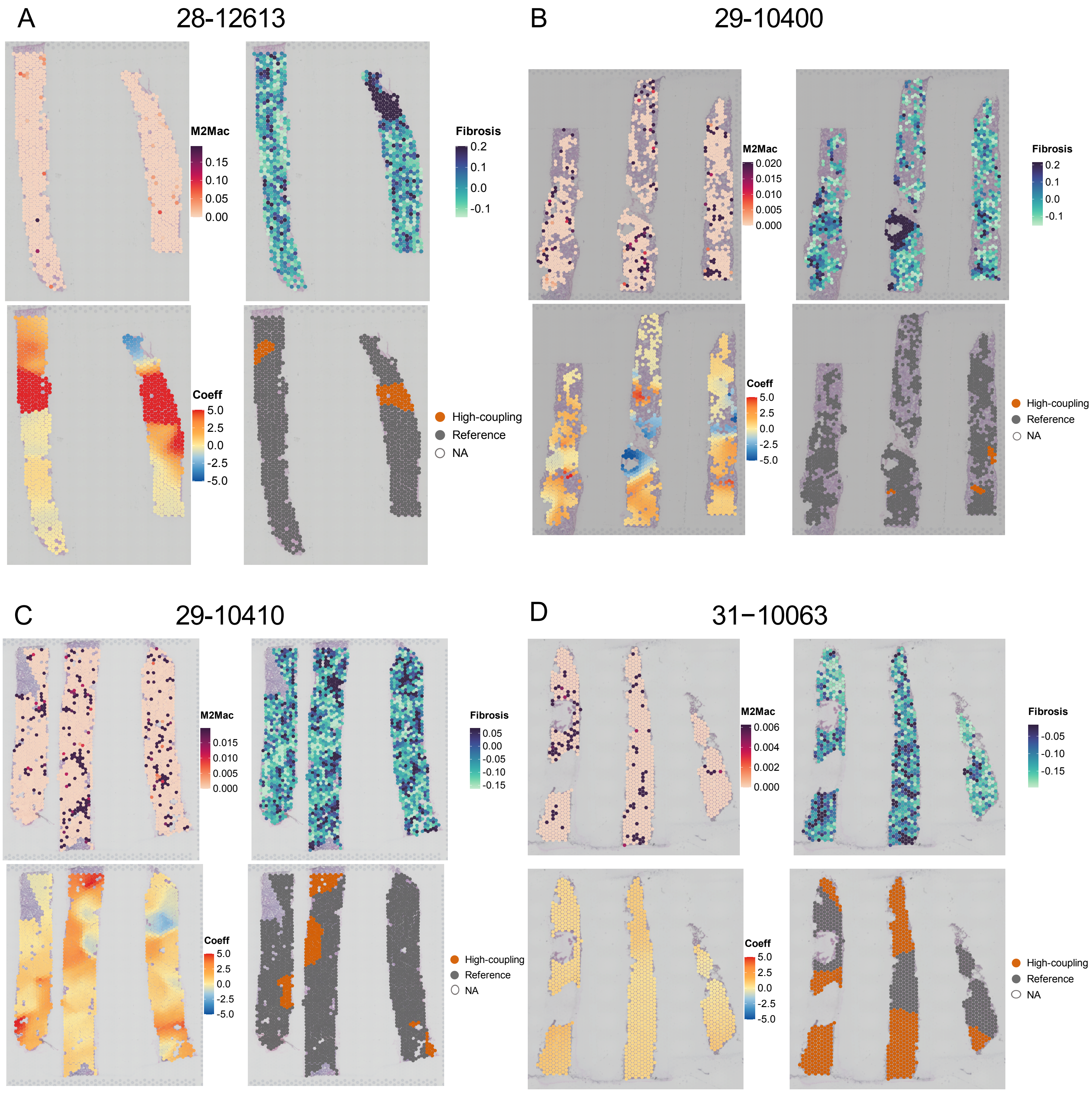

### Figure S5. GWR-based spatial analysis in KPMP HKD samples (part 2)

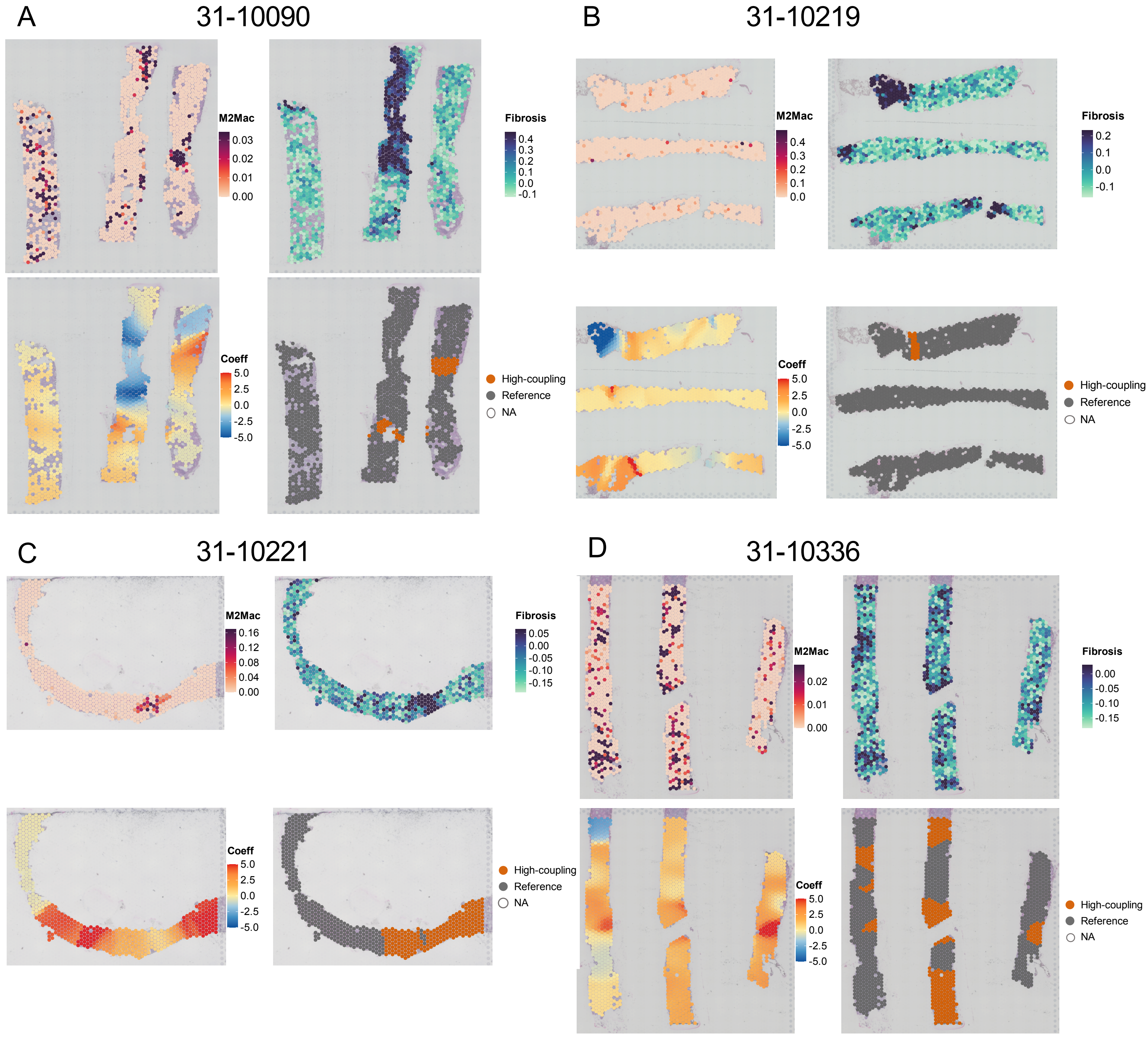
